# Decoding Odor Mixtures in the Dog Brain: An Awake fMRI Study

**DOI:** 10.1101/754374

**Authors:** Ashley Prichard, Raveena Chhibber, Jon King, Kate Athanassiades, Mark Spivak, Gregory S. Berns

## Abstract

In working and practical contexts, dogs rely upon their ability to discriminate a target odor from distracting odors and other sensory stimuli. Few studies have examined odor discrimination using non-behavioral methods or have approached odor discrimination from the dog’s perspective. Using awake fMRI in 18 dogs, we examined the neural mechanisms underlying odor discrimination between two odors and a mixture of the odors. Neural activation was measured during the presentation of a target odor (A) associated with a food reward, a distractor odor (B) associated with nothing, and a mixture of the two odors (A+B). Changes in neural activation during the presentations of the odor stimuli in individual dogs were measured over time within three regions known to be involved with odor processing: the caudate nucleus, the amygdala, and the olfactory bulbs. Average activation within the amygdala showed that dogs maximally differentiated between odor stimuli based on the stimulus-reward associations by the first run, while activation to the mixture (A+B) was most similar to the no-reward (B) stimulus. To identify the neural representation of odor mixtures in the dog brain, we used a random forest classifier to compare multilabel (elemental) vs. multiclass (configural) models. The multiclass model performed much better than the multilabel (weighted-F1 0.44 vs. 0.14), suggesting the odor mixture was processed configurally. Analysis of the subset of high-performing dogs based on their brain classification metrics revealed a network of olfactory information-carrying brain regions that included the amygdala, piriform cortex, and posterior cingulate. These results add further evidence for the configural processing of odor mixtures in dogs and suggest a novel way to identify high-performers based on brain classification metrics.

## 1. Introduction

For working purposes, trained dogs are generally considered the most practical and effective means of identifying target substances. In many cases, detection dogs are selectively bred for olfactory capabilities and behavioral traits that are correlated with their effectiveness in the field. Given their roles in national security and in detecting different diseases, hunting for pests, and tracking endangered species for conservation efforts, odor detection dogs remain in high demand (Bijland, Bomers, & Smulders, 2013; Cooper, Wang, & Singh, 2014; Davidson, Clark, Johnson, Waits, & Adams, 2014; Gadbois & Reeve, 2014). Research regarding dogs’ olfactory abilities typically focuses on the improvement of detection behaviors and trainability. Despite numerous behavioral studies, little is known about the way in which olfactory information is interpreted by the canine brain. Few studies on canine olfaction approach the topic from the canine’s point of view or without responses mediated by the dog’s handler. While behavior is a necessary measure of a working dog’s effectiveness, a dog’s behavior can be biased by unconscious cues given by their handler.

Large gaps remain in our understanding of how dogs process odors or discriminate between pure odors and their mixtures. For instance, it is unknown whether dogs search for the complete odor signature of a target substance or whether only some components serve as a target odor (Johnen, Heuwieser, & Fischer-Tenhagen, 2017). Despite substantial training on odor components, a dog’s behavioral responses to mixtures often cannot be predicted. This may be because the detection of individual substances within a mixture depends on chemical interactions between the different components. Given that most odor discrimination tests for dogs are behaviorally based and/or unstandardized, it is almost impossible to predict which components of an odor a particular dog uses to identify the target (Göth, McLean, & Trevelyan, 2003). For example, dogs that were trained to detect pure potassium chlorate failed to reliably detect potassium chlorate-based explosive mixtures (Lazarowski & Dorman, 2014). Whereas dogs trained on odor mixtures tend to perform better on detection tasks than when trained on pure odors (Hall & Wynne, 2018). These findings highlight the potential limitations of training dogs to detect a specific target odor to then indicate to the target when mixed with distractors (DeGreeff et al., 2017; Hayes, McGreevy, Forbes, Laing, & Stuetz, 2018). The way in which this information is interpreted by the canine brain also remains under-researched, but it is likely a complex and contextually dependent process (Berns, Brooks, & Spivak, 2015; Hayes et al., 2018; Prichard, Chhibber, Athanassiades, Spivak, & Berns, 2018; Siniscalchi, 2016). Considering that olfactory neuroanatomy is highly conserved among animals, studies of olfactory processing in dogs may also shed light on similar mechanisms in humans (Ache & Young, 2005).

The brain may have specialized representations for olfactory associations (Yeshurun, Lapid, Dudai, & Sobel, 2009). In humans, studies of odor perception typically rely on self-report measures and use suprathreshold odors that are easily detectible. Functional magnetic resonance imaging (fMRI) of an olfactory matching or identification task has demonstrated activation in the primary and secondary olfactory regions including: the piriform cortex, insula, amygdala, parahippocampal gyrus, caudate nucleus, inferior frontal gyrus, middle frontal gyrus, superior temporal gyrus, and cerebellum (Vedaei et al., 2017). Low level odors that go unnoticed by participants can also alter brain activation in the piriform cortex and thalamus (Lorig, 2012). Most of these studies have contributed to the identification of odor processing regions, but fewer have identified the regions’ roles during odor processing or learning during conditioning to odor stimuli. Regions that are thought to support conditioned associations to odors include the orbitofrontal and perirhinal cortices (Howard, Kahnt, & Gottfried, 2016; Qu, Kahnt, Cole, & Gottfried, 2016).

More recent studies of human olfactory perception have implemented machine learning strategies to decode odor representation within the brain. FMRI decoding methods can reveal regions important for coding valence, expected outcomes, or stimulus identity. Machine learning approaches, such as multi-voxel pattern analysis (MVPA) or representational similarity analysis (RSA), identify patterns of activation from regions that might not show a change in mean activation with univariate measures (Haxby, Connolly, & Guntupalli, 2014; Kahnt, 2018; Kahnt, Park, Haynes, & Tobler, 2014). In another study using RSA suggested that the spatial and temporal pattern of activation within the amygdala codes for odor valence (Jin, Zelano, Gottfried, & Mohanty, 2015).

While these studies report similar regions important for odor discrimination identified in traditional univariate fMRI analyses, the relationship of odor mixtures to the brain’s representation of odor components remains unknown (Howard & Gottfried, 2014; Howard, Gottfried, Tobler, & Kahnt, 2015; Howard et al., 2016). Odor mixtures may be represented in the brain based on their components (elemental) or may be perceived as configural, creating an odor concept (Thomas-Danguin et al., 2014). FMRI studies on human perception of odor mixtures shows that activation in the insula increases when the participant experiences the mixture containing the target odor, even when participants report that they are unable to distinguish the mixture with and without the target (Hummel, Olgun, Gerber, Huchel, & Frasnelli, 2013). However, other regions identified in this study included voxel sizes that would not pass whole brain corrections for multiple comparisons, requiring further study to confirm these brain regions’ roles in the perception of odor mixtures. Despite the need for the study of human olfactory perception at the neural level, no research has yet investigated similar considerations in the dog (Hayes et al., 2018).

Studies of canine cognition using fMRI are becoming more common, including the adaptation of human experimental paradigms and analyses. With appropriate selection and training, dogs can be willing participants in fMRI and show little anxiety in the testing environment as it is similar to their shared environment with humans. Due to domestication, dogs are also more likely attuned to stimuli relevant to humans as opposed to stimuli salient to other model species. Since 2012, dog fMRI has revealed some of the conserved neural mechanisms underlying perception across species (Berns, Brooks, & Spivak, 2012). Dogs have a region for processing both human and dog faces similar to that of primates (Cuaya, Hernandez-Perez, & Concha, 2016; Dilks et al., 2015). Dogs show differential activation in the reward processing regions of the brain such as the caudate nucleus to social or food rewards (Cook, Prichard, Spivak, & Berns, 2016). And dogs show higher activation in the amygdala and caudate to odors associated with familiar humans and dogs than to odors of strangers (Berns et al., 2015). Canine fMRI studies have also revealed neural biases for stimulus modalities, suggesting that dogs learn visual and odor stimuli at a faster rate than verbal stimuli, and that differences in activation are most evident in the amygdala and caudate (Prichard, Chhibber, et al., 2018). Finally, MVPA analysis of dog fMRI data revealed that dogs and humans have similar brain regions for the representation of semantic knowledge in the form of words associated with objects (Prichard, Cook, Spivak, Chhibber, & Berns, 2018). Together, these studies suggest that dogs are not only willing fMRI participants, but that the existence of functionally similar brain regions shared by dogs and humans make them an appropriate model species for further research.

To examine the neural mechanisms underlying a dog’s classification of odor mixtures, we measured the fMRI response to two previously trained odors (one associated with reward and one not) (Prichard, Chhibber, et al., 2018) and to a mixture of the two odors. First, we used univariate analyses on mean activation levels within the olfactory bulb, amygdala, and caudate nucleus to determine whether the mixture was more similar to the pure reward or no-reward odors. Second, we used a random forest classifier (RFC) for: a) whole-brain decoding of odor identity; b) determination of whether a mixture is processed elementally or configurally; and c) identification of additional regions for odor classification in the dog’s brain.

## 2. MATERIALS AND METHODS

### 2.1 Participants

Participants were 18 pet dogs volunteered by their Atlanta owners for fMRI training and fMRI studies (Berns, Brooks, & Spivak, 2013; Berns et al., 2012; Berns & Cook, 2016; Cook et al., 2016). All dogs had previously completed four or more awake fMRI scans, including previous training on the two odors used in the current study (Prichard, Chhibber, et al., 2018). No physical or chemical restraint was implemented. The study utilized odor stimuli that each dog had previously experienced within the scanner environment. This study was performed in accordance with the recommendations in the Guide for the Care and Use of Laboratory Animals of the National Institutes of Health. The study was approved by the Emory University IACUC (Protocols DAR-4000079-ENTPR-A and PROTO201700572), and all owners gave written consent for their dog’s participation in the study.

### 2.2 Stimuli

Olfactory stimuli were aqueous solutions of isoamyl acetate (IA), hexanol (Hex), and a mixture of the two calculated to result in approximately 5 ppm in the headspace of the container. Partial vapor pressures were calculated based on the molecular weight and reported vapor pressures of 4 mmHg and 0.9 mmHg respectively, obtained from PubChem (pubchem.ncbi.nlm.nih.gov). The odorants were miscible with water and the partial pressure of the odorant was the product of the pure odorant vapor pressure and the mole fraction of the odorant. The final dilutions in water were 0.12 mL/L for IA, 0.44 mL/L for Hex.

Odorants were delivered using an MRI-compatible olfactometer used in a previous study and similar to those constructed for human olfactory imaging studies (Bestgen et al., 2016; Lowen & Lukas, 2006; Prichard, Chhibber, et al., 2018; Sezille et al., 2013; Sommer et al., 2012; Toledano et al., 2012; Vigouroux, Bertrand, Farget, Plailly, & Royet, 2005). Briefly, odorants were delivered using a continuous stream of air from an aquarium grade air pump (EcoPlus Commercial Air Pump 1030 GPH) through a Drierite filter (drierite.com), and through a 4-way plastic splitter to three plastic 100 mL jars containing 50 ml of odorant solutions and one jar containing 50 ml of water to serve as a control. Each solution mixed with a continuous air stream. The experimenter used plastic valves to control directional flow of odorized air through 10’ of 1/8” ID Teflon tube, where the mixture (air dilution of the odorant) exited a PVC tube with a 1” diameter opening positioned in the MRI bore 12” from the dog’s snout (Fig. 1). The fourth tube carrying air from the control jar remained open throughout the presentations of odorized air, maintaining a steady air stream presented to the dog and assisting in the clearing of lingering odor within the magnet bore.

**Figure 1.**
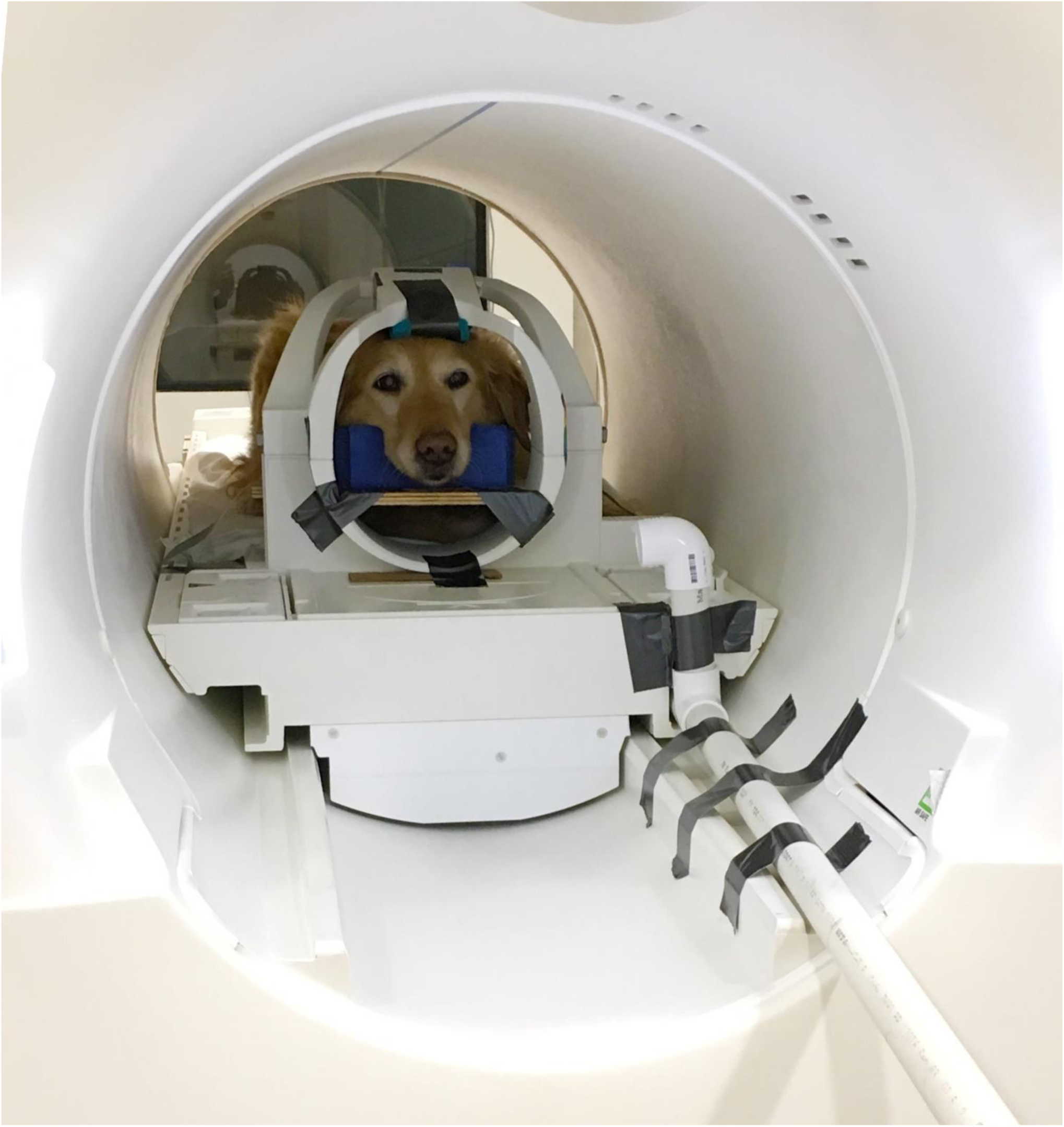
Experimental design with odor stimuli. Three odor stimuli were repeatedly presented during a scan session. One stimulus was associated with food (Reward), while the No Reward and Mixture stimuli were associated with nothing. Presentation of odorants to dog in MRI bore via experimenter-controlled olfactometer during scan session. The owner remained in front of the dog.

### 2.3 Experimental Design

Dogs entered and stationed themselves in custom chin rests in the scanner bore. All scans took place in the presence of the dog’s primary owner, who stood throughout the scan at the opening of the magnet bore, directly in front of the dogs, and delivered all rewards (hot dogs) to the dog. The owner was present to minimize any anxiety that the dog may experience due to separation, consistent with studies involving pets or human infants. An experimenter was stationed next to the owner, out of view of the dog. The experimenter controlled the timing and presentation of odor stimuli to the dogs via a four-button MRI-compatible button box. Onset of each stimulus was timestamped by the simultaneous press of the button box with the opening of the appropriate valve. Manual control of the stimuli by the experimenter was necessary, as opposed to a scripted presentation, because of the variable time it takes dogs to consume food rewards.

In a previous study, dogs were semi-randomly assigned IA or Hex as the reward stimulus such that roughly half of the dogs were assigned to each group (see Table 1) (Prichard, Chhibber, et al., 2018). In the current study, the same dogs were presented with the two previously trained odors, as well as a mixture of the two. An event-based design was used, consisting of reward, no-reward, and mixture trial types. On reward trials, the odor stimulus was presented for a fixed duration, which was followed by the delivery of a food reward. During no-reward trials and mixture trials, the no-reward or mixture odor stimuli were presented for the same fixed duration and were followed by nothing. Each dog received the same trial sequence. For each trial type, dogs were presented an odor for an initial 3.6s during a span of 7.2 s, followed by a reward (hot dog) or nothing, with a 9.6 s inter trial interval between odor presentations.

**Table 1.**
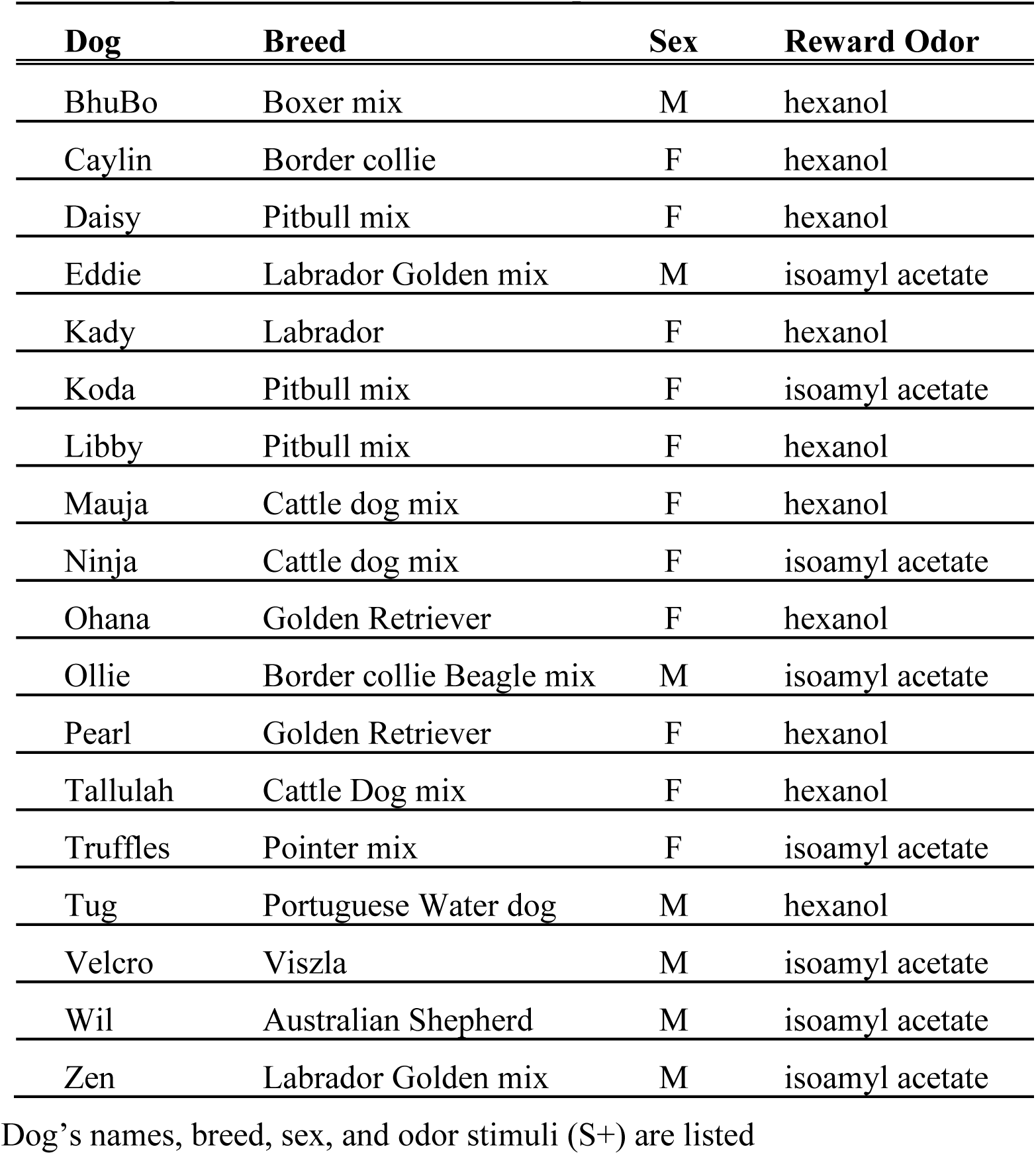
Dogs (N=18) and odor stimulus paired with reward.

| Dog | Breed | Sex | Reward Odor |
| --- | --- | --- | --- |
| BhuBo | Boxer mix | M | hexanol |
| Caylin | Border collie | F | hexanol |
| Daisy | Pitbull mix | F | hexanol |
| Eddie | Labrador Golden mix | M | isoamyl acetate |
| Kady | Labrador | F | hexanol |
| Koda | Pitbull mix | F | isoamyl acetate |
| Libby | Pitbull mix | F | hexanol |
| Mauja | Cattle dog mix | F | hexanol |
| Ninja | Cattle dog mix | F | isoamyl acetate |
| Ohana | Golden Retriever | F | hexanol |
| Ollie | Border collie Beagle mix | M | isoamyl acetate |
| Pearl | Golden Retriever | F | hexanol |
| Tallulah | Cattle Dog mix | F | hexanol |
| Truffles | Pointer mix | F | isoamyl acetate |
| Tug | Portuguese Water dog | M | hexanol |
| Velcro | Viszla | M | isoamyl acetate |
| Wil | Australian Shepherd | M | isoamyl acetate |
| Zen | Labrador Golden mix | M | isoamyl acetate |

Dog's names, breed, sex, and odor stimuli (S+) are listed

Each scan session consisted of 4 runs, lasting approximately 9 minutes per run. Each run consisted of 22 trials (∼8 reward, ∼8 no-reward, ∼5 mixture) with a semi-randomized presentation order, for a total of 88 trials per scan session. Twenty-two mixture trials were included to serve as a sufficient number of probe trials for fMRI analyses while minimizing mixture-outcome associations. No trial type was repeated more than 3 times sequentially, as dogs could habituate to the stimulus. Following each run, dogs would exit the scanner and relax, drink water, or stay in the scanner to complete the next run.

Scanning was conducted with a Siemens 3 T Trio whole-body scanner using procedures described previously (Berns et al., 2013; Berns et al., 2012). During the first of the four runs, a T2-weighted structural image of the whole brain was acquired using a turbo spin-echo sequence (25-36 2mm slices, TR = 3940 ms, TE = 8.9 ms, flip angle = 131°, 26 echo trains, 128 × 128 matrix, FOV = 192 mm). The functional scans used a single-shot echo-planar imaging (EPI) sequence to acquire volumes of 22 sequential 2.5 mm slices with a 20% gap (TE = 25 ms, TR = 1200 ms, flip angle = 70°, 64 × 64 matrix, 3 mm in-plane voxel size, FOV = 192 mm). Slices were oriented dorsally to the dog’s brain (coronal to the magnet, as in the sphinx position the dogs’ heads were positioned 90 degrees from the prone human orientation) with the phase-encoding direction right-to-left. Sequential slices were used to minimize between-plane offsets from participant movement, while the 20% slice gap minimized the “crosstalk” that can occur with sequential scan sequences. Four runs of up to 400 functional volumes were acquired for each subject, with each run lasting about 9 minutes.

### 2.4 Analyses

#### 2.4.1 Preprocessing

Preprocessing of the fMRI data included motion correction, censoring, and normalization using AFNI (NIH) and its associated functions. Two-pass, six-parameter rigid-body motion correction was used based on a hand-selected reference volume for each dog that corresponded to their average position within the magnet bore across runs. Aggressive censoring removed unusable volumes from the fMRI time sequence because dogs can move between trials, when smelling an odor, and when consuming rewards. Data were censored when estimated motion was greater than 1 mm displacement scan-to-scan and based on outlier voxel signal intensities greater than 0.1 percent signal change from scan-to-scan. Smoothing, normalization, and motion correction parameters were identical to those described in previous studies (Prichard, Chhibber, et al., 2018). EPI images were smoothed and normalized to %-signal change with 3dmerge using a 6mm kernel at full-width half-maximum. The Advanced Normalization Tools (ANTs) software was used to register the mean of the motion-corrected functional images (Avants et al., 2011) to the individual dog’s structural image.

#### 2.4.2 Region of Interest (ROI) Analysis

Each subject’s motion-corrected, censored, smoothed images were analyzed within a general linear model (GLM) for each voxel in the brain using 3dDeconvolve (part of the AFNI suite). Motion time courses were generated through motion correction, and constant, linear, quadratic, cubic, and quartic drift terms were included as nuisance regressors. Drift terms were included for each run to account for baseline shifts between runs as well as slow drifts unrelated to the experiment. Task related regressors for each experiment were modeled using AFNI’s dmUBLOCK and stim_times_IM functions and were as follows: (1) reward stimulus, (2) no-reward stimulus, 3) mixture stimulus. The function created a column in the design matrix for each of the 88 trials, allowing for the estimation of a beta value for each trial. Trials with beta values greater than an absolute three percent signal change were removed prior to analyses as described in Prichard et al. (2018) as these were assumed to be beyond the physiologic range of the BOLD signal and possibly the result of spin-history effects and spurious levels of activation unrelated to the experiment.

Anatomical ROIs were selected based on imaging results in canine brain areas previously observed to be responsive to olfactory stimuli (Berns et al., 2015; Jia et al., 2014). Anatomical ROIs of the left and right caudate nuclei, the left and right amygdala, and the olfactory bulbs were defined structurally using each dog’s T2-weighted structural image of the whole brain (Fig. 2). Beta values for each presentation of reward stimuli (33 trials), no-reward stimuli (33 trials), and mixture stimuli (22 trials) were extracted from and averaged over the ROIs in the left and right hemispheres. For each ROI (amygdala, caudate, olfactory bulb), we used the mixed-model procedure in SPSS 24 (IBM) with fixed-effects for the intercept, run number, type (reward, no-reward, mixture), and hemisphere (left or right), identity covariance structure, and maximum-likelihood estimation. Run was modeled as a fixed effect, making no assumptions about the time course. As hemisphere did not account for a significant amount of variance, data were collapsed across hemispheres and analyses removed hemisphere as a factor.

**Figure 2.**
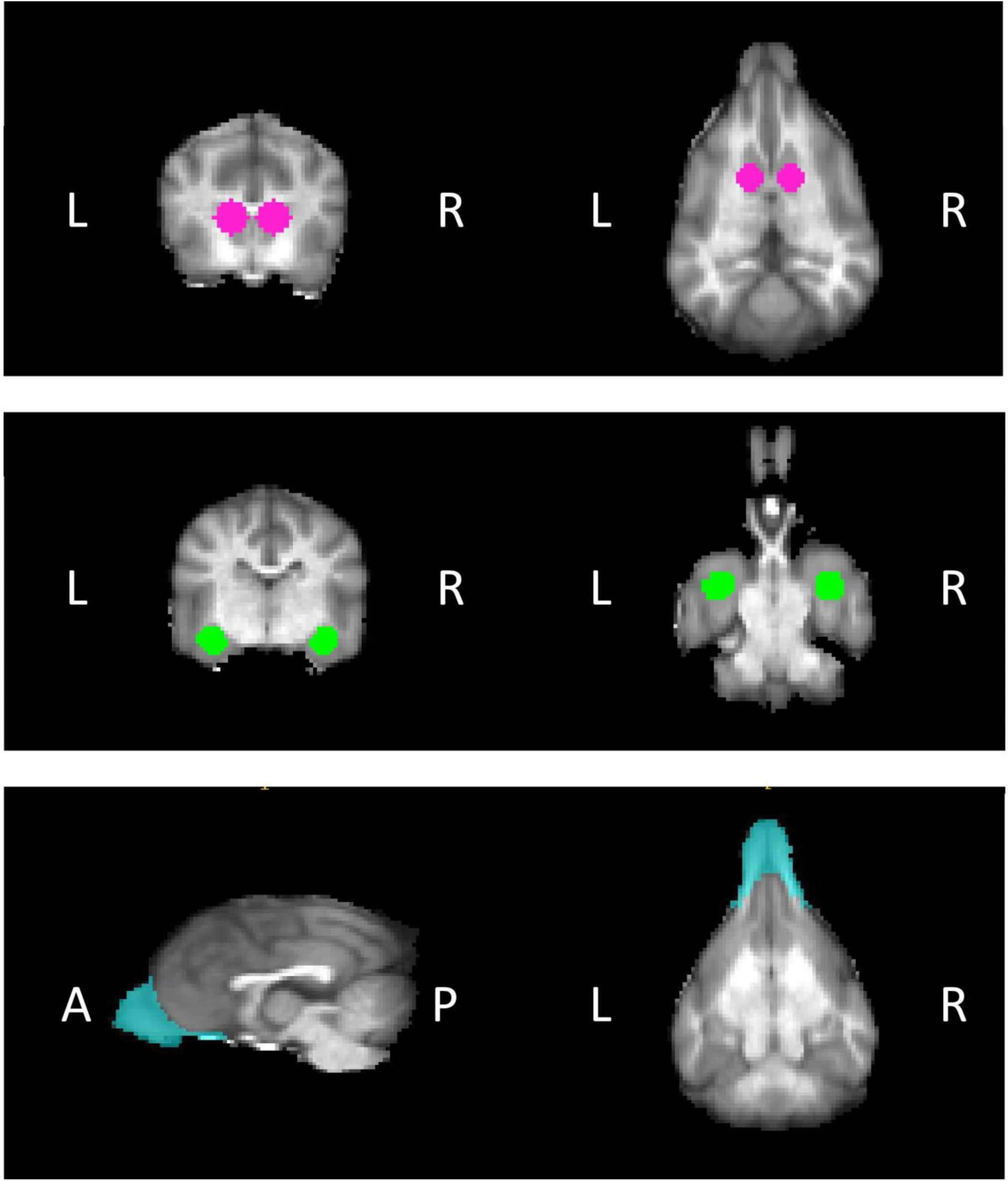
Regions of interest (ROIs). ROIs were drawn in individual anatomical space, example ROIs shown in template space here in transverse and dorsal views. **A)** Caudate nuclei have been shown to differentially respond to odor stimuli associated with reward and no-reward. **B)** Amygdalae have shown differential responding to odor stimuli associated with reward and no-reward, as well as arousal. **C)** Olfactory bulbs including olfactory bulbs respond to odor stimuli. ROI is shown here in sagittal and dorsal views in template space.

#### 2.4.3 Multivariate Decoding

For this exploratory analysis, our aim was to identify regions in the dog brain that contribute to the classification of odor stimuli outside of those identified in the univariate analysis. Univariate analyses may answer the question if odor mixtures result in differences in regional brain activity, but multivariate methods are required if the identity of odors is distributed in patterns of neural activity. The primary question was whether dogs treat odor mixtures as elemental or configural. As in previous decoding human fMRI studies, we used scikit-learn’s random forest classifier (RFC). RFC has previously demonstrated robust performance on human fMRI data and has the ability to handle complex biological data (Lebedev et al., 2014). RFCs generally perform better than most linear classifiers and require less parameter tuning (Chollet, 2018). An RFC also allows for mapping of feature importance in the brain without resorting to searchlight analyses. Thus, in addition to generating whole-brain classification metrics, the relative importance of individual regions to the classification can be obtained.

The volumes from the current study were concatenated with data from the previous study in which dogs were presented odors associated with reward and no reward in a classical conditioning paradigm (Prichard, Chhibber, et al., 2018), yielding a total of 176 separate odor trials. As described in the above GLM, preprocessing included censoring of the unsmoothed volumes for motion and outliers. Using AFNI’s 3dDeconvolve stim_times_IM function, we generated a whole-brain model of trial-by-trial beta estimates for each trial type (reward, no reward, and mixture). The anatomical masks from the ROI analysis described above were used to extract average beta values from the left and right caudate for each trial. As in the univariate analysis, trials with beta values greater than |3 %| were removed prior to further analyses. Using AFNI’s 3dmerge tool, the remaining whole brain volumes were smoothed with a kernel of 6 mm to improve signal-to-noise ratios. The whole brain volumes were used as input for the classifiers below. To reformat the imaging data for use in the sklearn environment, the volumes were masked and reshaped using nilearn’s NiftiMasker class and split into training and testing sets using the python library *pandas*.

Two different models were tested: elemental and configural. For the elemental model, trials were coded using a 2-bit vector with bits for odor A and odor B. In this scheme, trials with the two pure odorants were coded as [1 0] and [0 1] while the mixture was coded as [1 1]. In contrast, the configural model assumed that the mixture was a distinct class and was coded as such. Here, the classes were simply A, B, and C. The primary difference between these two models was multilabel vs. multiclass.

For both models, the RFC was instantiated in each dog by making 100 forests, each forest consisting of 100 trees with a max_depth of 5, min_samples_split of .25, bootstrapping as true, and max_features as log2. We used 100 forests of 100 trees to ensure that all volumes served as samples. A max_depth of 5, min_samples_split of .25 and max_features of log2 were included to prevent overfitting to the training set. Each dog’s data was split into odd and even runs (2-fold split) for training and testing. For training each forest, an equal number of exemplars from each class was randomly selected. Unselected trials were added to the test set. For each trial of the test set, the classifier predicted whether the stimulus presented was reward, no reward, or mixture. From this, we calculated the confusion matrix for each dog, aggregating over the 100 forests. The primary metrics obtained were recall, precision, and the F1-score (a weighted average of recall and precision).

Each forest also produced a map of feature importances. Briefly, the feature importance is a value scaled between 0 and 1 that reflects how informative a voxel i.e. a larger feature importance corresponds to a voxel that is more informative in making the final predictions. Higher feature importances are driven by either voxels that increase accuracy drastically, or by voxels that are present in many trees within a forest. Sklearn’s RFC feature_importances_ method returned feature importances for each voxel that were subsequently back-mapped into each individual dog’s functional space, generating one map per forest. All 100 maps for each dog were averaged to assess which brain regions contributed to the classification reward, no reward, and mixture. Mean images for each dog were spatially normalized to template space using The Advanced Normalization Tools (ANTs) software (Avants et al., 2011; Datta et al., 2012).

To determine the significance of both the confusion matrices and feature importance maps, we followed the permutation approach outlined by Stelzer et al. (2013) and which we used previously to identify the significance of regions for language processing in dogs (Prichard, Cook, et al., 2018; Stelzer, Chen, & Turner, 2013). For each dog, a random number was appended to the data labels for each trial to reorder the labels and create a permuted list of labels, while the timeseries of fMRI volumes remained unchanged. The RFC was trained and tested on this set of permuted labels and the fMRI volumes 100 times, outputting a confusion matrix and a map of feature importances for each forest. As we did with our real dataset above, we then averaged across these 100 forests, creating one confusion matrix and one mean image per set of permuted labels. We repeated this procedure 100 times to create a distribution of confusion matrices and feature importance maps for each dog.

For each confusion matrix, we computed the weighted F1 score. This allowed us to calculate the cumulative distribution of F1 scores for the permuted data, which then allowed an estimation of the significance of the actual F1 score for the real data. As we were interested in identifying additional brain regions involved in the identification of odors, we included those dogs whose whole-brain classifier performed substantially above chance. Only dogs who had a real F1 greater than the 90^th^ percentile of the null distribution were used to create a group feature importance map.

To simulate the group image across dogs, we randomly selected one mean permuted image per dog, normalized that mean image to template space, and averaged across the dogs comprising the group map – i.e. those dogs whose F1 was greater than the 90^th^ percentile of their null distribution. This random selection and normalization were repeated 10,000 times. Because each voxel in the brain may have a different distribution given its location in the brain, we did not assume a canonical distribution across all voxels and opted to make a voxel-wise distribution. For each voxel in the brain, we created the distribution from the 10,000 noise maps and determined the values for *p* = 0.005. This map of thresholds was applied to the mean feature importance map created from the real data and to each of the 10,000 noise maps (Fig. 3). To determine the significance of any clusters found after thresholding at the voxel-wise level, we created a distribution of cluster sizes found in the thresholded 10,000 noise maps.

**Figure 3.**
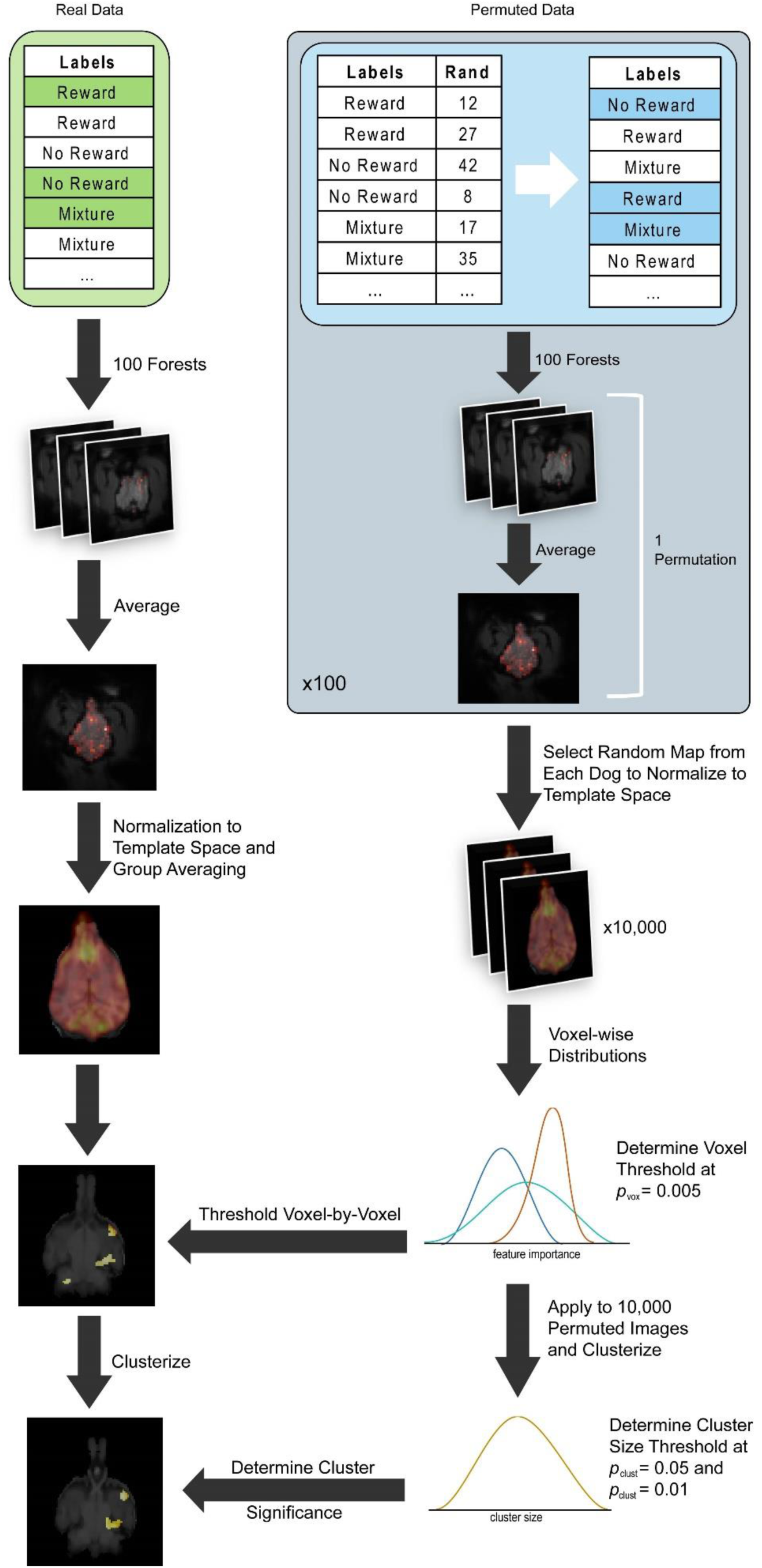
Schematic diagram of MVPA methods. A random forest classifier (RFC) was trained on a balanced subset of the real data, outputting a map of voxels important for classification. We repeated this process 100 times to ensure that all samples were used at least one. The maps from these 100 repetitions were averaged, normalized to group space, and thresholded at a voxel and cluster level to create the final image. To determine the voxel and cluster-level thresholds, we created random data by permuting the data labels associated with each volume, then trained as described above for the real data, which constituted one permutation. The data were permuted 100 times and one map was selected at random to transform into group space. We generated 10,000 random group maps, created a voxel-by-voxel distribution and a cluster distribution, which were then applied to the image generated by the real data.

## 3. RESULTS

### 3.1 Univariate

Changes in neural activation during the presentations of the odor stimuli in individual dogs were measured over time within the three ROIs known to be involved with odor processing. Using the mixed-model procedure in SPSS 24 (IBM) we found neural evidence for differentiation of the three odor stimuli across all ROIs (*p* = 0.004), which varied significantly by Odor Type (*p* < 0.001). There was a significant interaction between Odor Type x Run (*p* = 0.031), suggesting the magnitude of the effect changed over time.

As there was a main effect of ROI, we used post-hoc analyses to examine whether these differences remained when segregated by ROI (Table 2 & Fig. 4). In the caudate, we found a significant interaction between Odor Type x Run (*p* = 0.019) (Fig. 5A), but no main effect of Odor Type or Run. More robust evidence for the differentiation between odor stimuli was evident in the amygdala for Odor Type (*p* < 0.0001) (Fig. 5B), suggesting that the odor-outcome associations were reinstated from the previous study. There was also an Odor Type x Run interaction, suggesting a difference in the temporal pattern between odor types (*p* = 0.028). Similar to human olfaction studies, we found initial evidence for the differentiation of Odor Type in the olfactory bulbs (*p* = 0.029) (Fig. 5C).

**Table 2.**
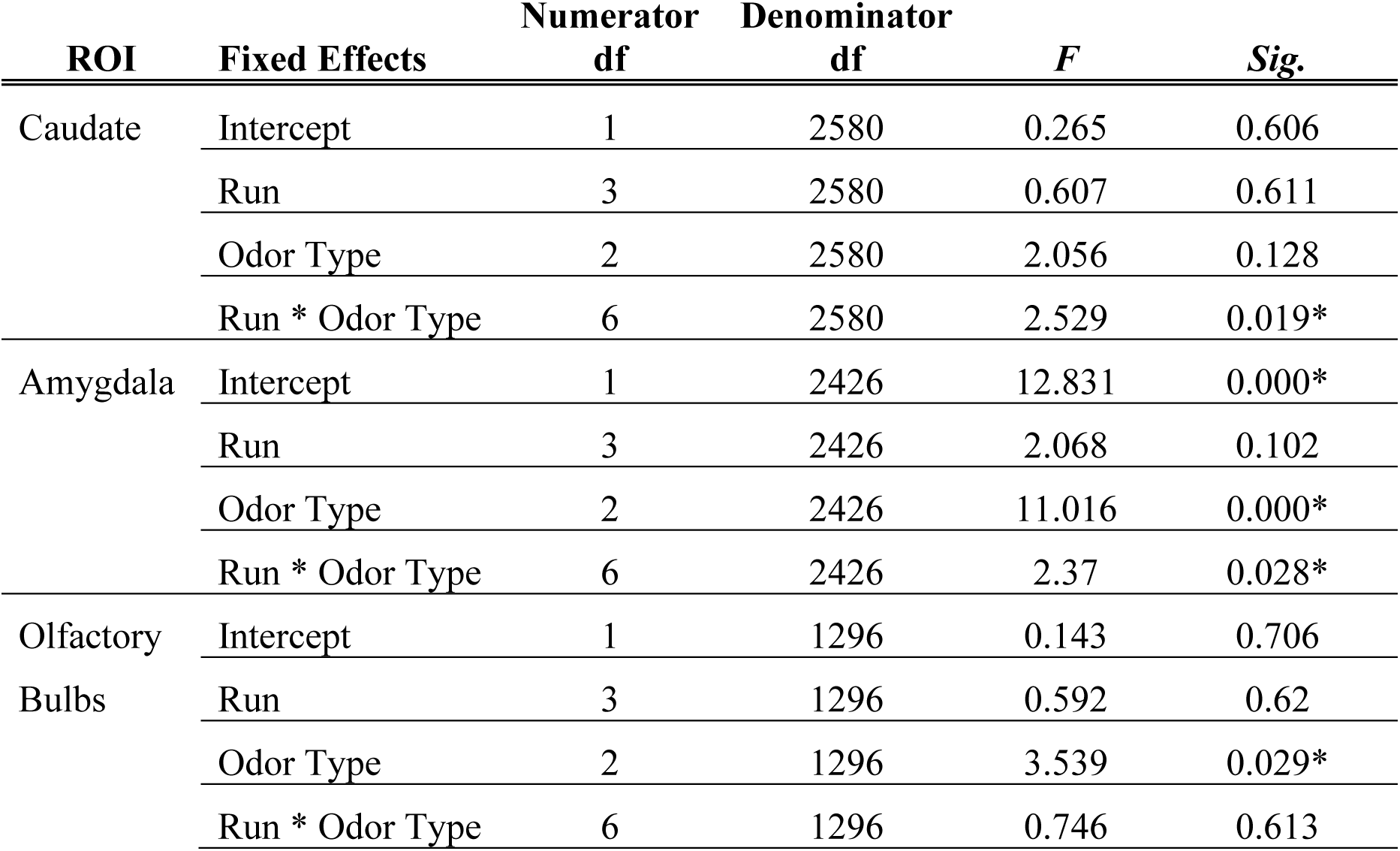
Model results for Odor Type, Run, and ROI. Asterisks denote significant results.

**Figure 4.**
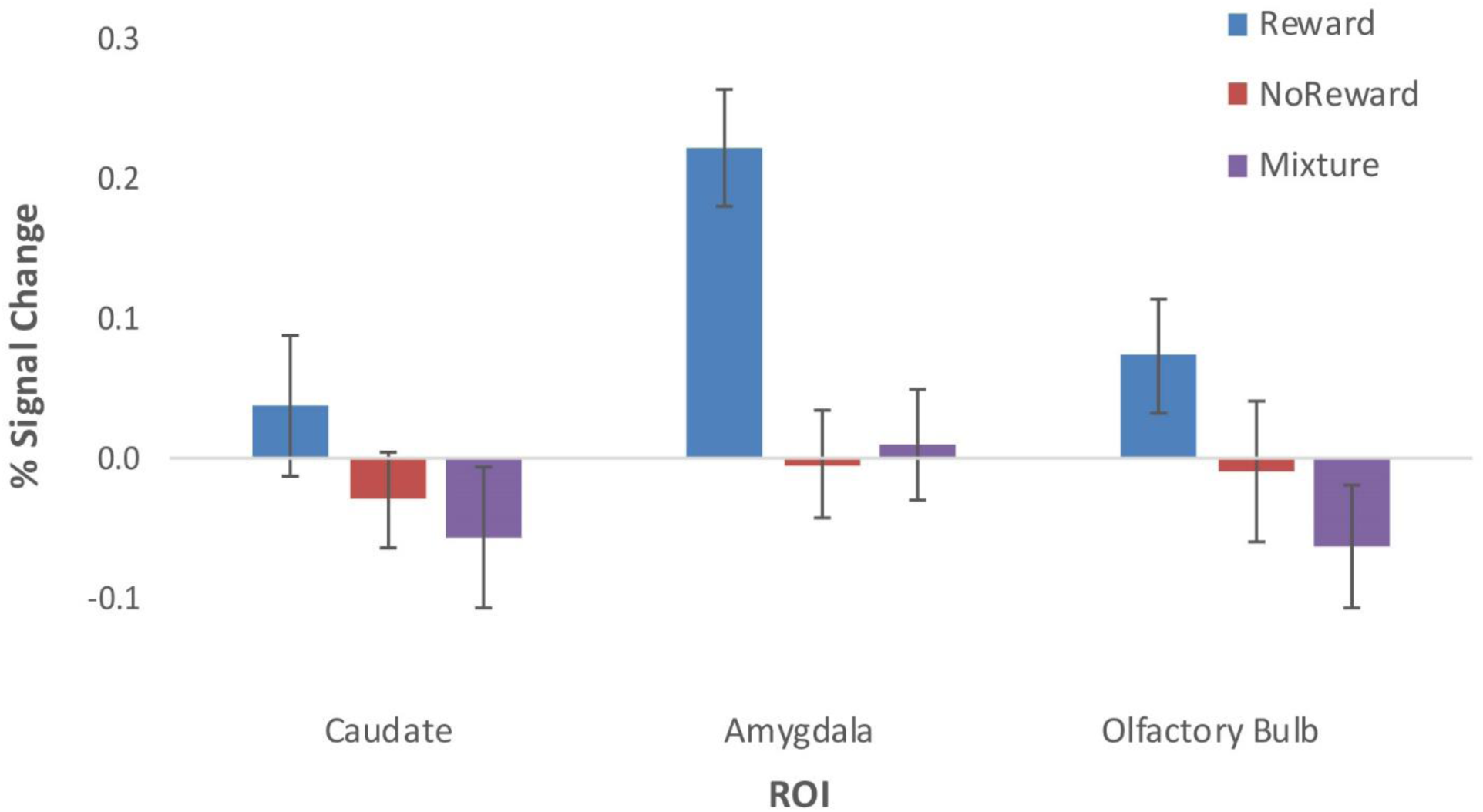
Percent signal change by ROI for odor stimuli. Mean values of odorant responses across dogs by ROI and by trial type are plotted relative to the implicit baseline (*blue* = reward, *red* = no reward, *purple* = mixture of reward and no reward). Error bars denote the standard error of the mean across dogs. Averaged beta values in the caudate did not show significant differentiation between odorants. The amygdala showed marked differentiation between odor stimuli, with the greatest activation to odor stimuli associated with reward. The olfactory bulbs followed a similar pattern of activation to the caudate. Across all ROIs, the neural activation to the mixture of odors was most similar to the neural response during the presentation of the no reward odor.

**Figure 5.**
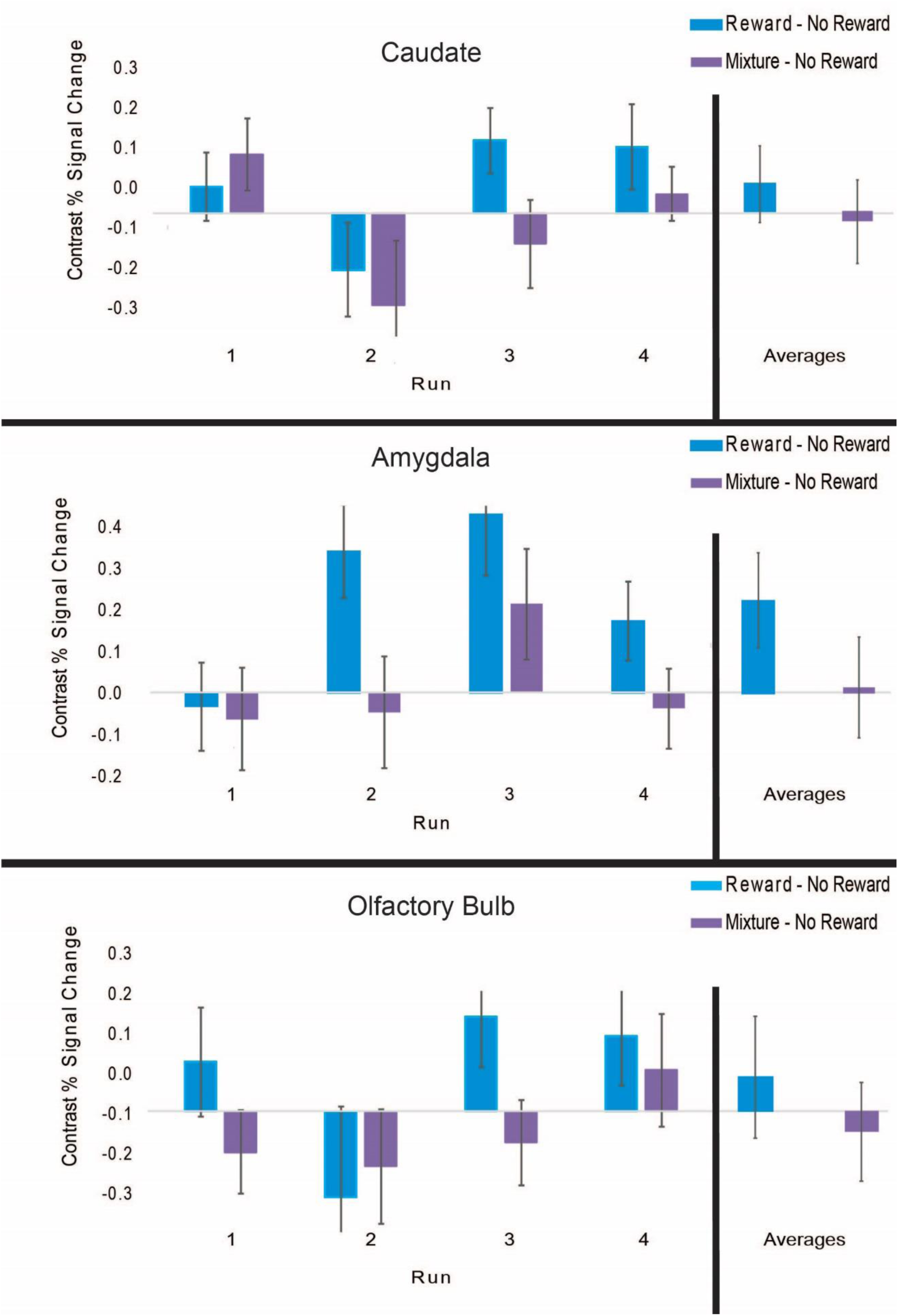
Percent signal change by ROI for reward and mixture odors relative to no reward odor. Mean values across dogs are plotted for each run (*blue* = Reward— No Reward, *purple* = Mixture— No Reward) and averages across all runs (*right*). Error bars denote the standard error of the mean across dogs. There were main effects of odor type across all ROIs (*p* = 0.004), which were significantly different by odor type (*p* < 0.001). There was a significant interaction ROI and Run (*p* = 0.031), suggesting the magnitude of the effect changed over time. **A)** Averaged beta values in the caudate show a significant interaction between Run and Odor Type (*p* = 0.036). **B)** Averaged beta values in the amygdala show significant effects of Odor Type (*p* = 0.001). **C)** Following corrections for multiple comparisons, activations in the olfactory bulbs were not significantly different.

In sum, the differences in neural activation across regions of the olfactory pathway show that dogs formed odor stimulus-reward associations. Though the differentiation between the three odor stimuli was most pronounced in the amygdala, similarity in activation between the no reward and mixture stimuli across all three ROIs suggested that the mixture was most like the no reward stimulus. However, when we tested whether the sum of activations to reward and no reward odors was the same as the activation to the mixture, we found significant differences in the amygdala, such that the sum of activations was greater than activation to mixture (*t*(17) = 3.28, *p* = 0.004). This suggests that mixture was, in fact, processed differently than the simple sum of its components. To further test this theory, multivariate decoding was performed.

### 3.2 Multivariate Decoding

Based on the weighted-F1 score, the multiclass model performed much better than the multilabel model (F1: 0.44 vs. 0.14) (Table 3). The multiclass model had an average recall of 0.40, which was better than the chance value of 0.33, while the multilabel model had very poor recall (0.19), in effect, predicting most stimuli as the mixture, including the pure odorants. Using the permuted data as a reference null distribution of F1 scores, we determined that the real data from 8 dogs passed the 90^th^ percentile (Bhubo, Caylin, Eddie, Kady, Koda, Ohana, Wil, and Zen). These dogs were then used to construct the whole brain map of informative voxels.

**Table 3.** Performance of multiclass and multilabel models.

|  |  | Precision | Recall | F1 |
| --- | --- | --- | --- | --- |
| Multiclass | Reward | 0.59 | 0.33 | 0.43 |
|  | No Reward | 0.55 | 0.46 | 0.50 |
|  | Mixture | 0.15 | 0.45 | 0.22 |
|  | Weighted Average | 0.52 | 0.40 | 0.44 |
| Multilabel | Reward | 0.66 | 0.03 | 0.06 |
|  | No Reward | 0.67 | 0.08 | 0.14 |
|  | Mixture | 0.12 | 0.91 | 0.21 |
|  | Weighted Average | 0.60 | 0.19 | 0.14 |

For these eight dogs, the feature importances of their multiclass models were backprojected into their brains, transformed to the atlas space, and then averaged. Only those voxels that passed the individual significance of *p* = 0.005 were used. Across these eight dogs, clusters with more than 2 voxels were used to create a cumulative distribution of possible cluster sizes. A cluster size of 98 voxels corresponded to *p* = 0.001. At this voxel and cluster threshold, three clusters were identified (Fig. 6). Two clusters surrounded the amygdala – one rostrally and one caudally. The third cluster was located in the posterior cingulate.

**Figure 6.**
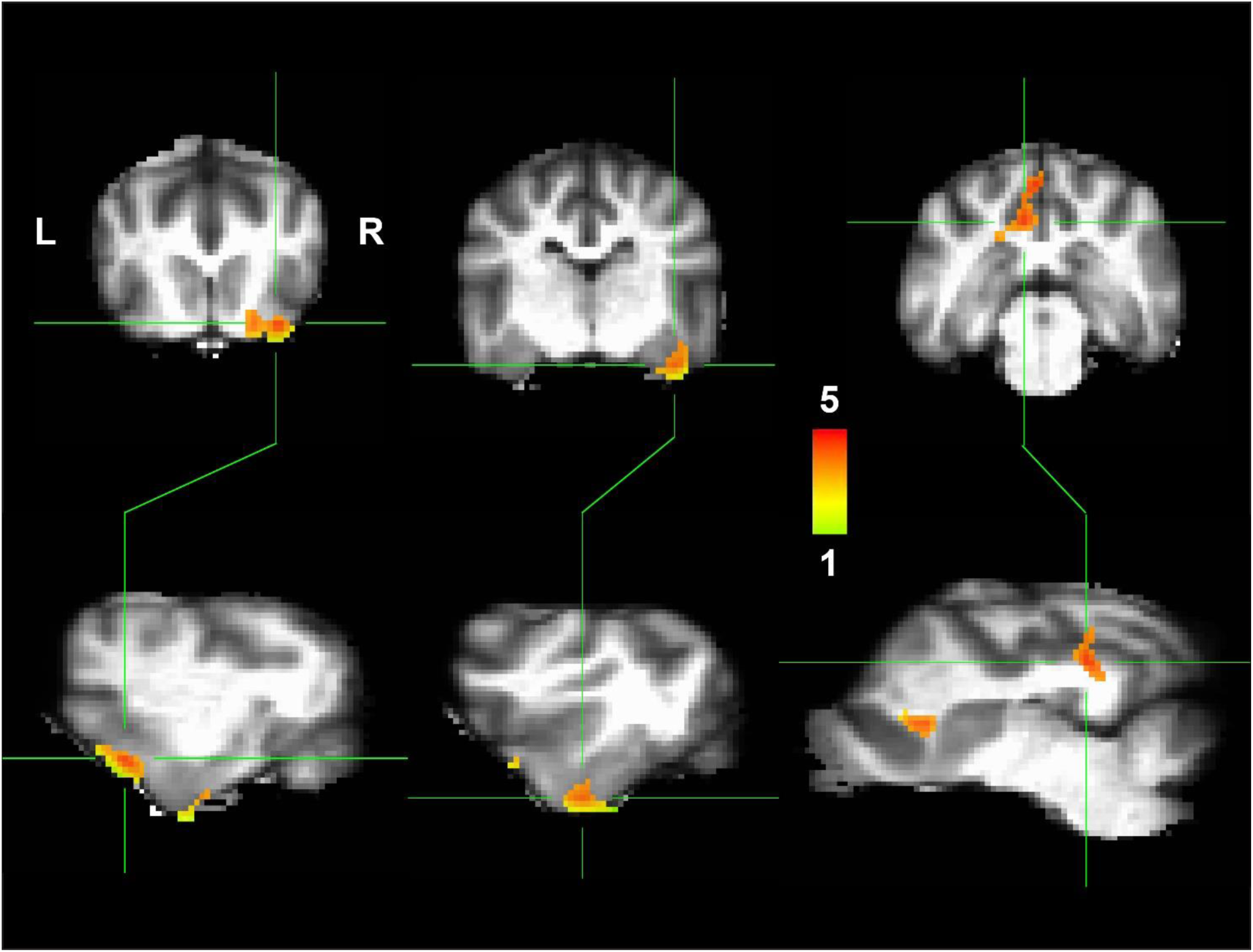
Clusters of informative voxels for multiclass random forest classifier. Three clusters were identified in the 8 dogs whose whole-brain classifier performed at the 90^th^ percentile of a null distribution. Two clusters bracketed the amygdala (*left* and *middle*) while the third cluster was located in the posterior cingulate (*right*). Voxel and cluster level significance is *p* = 0.005 and *p* = 0.001 respectively. Color indicates feature importance in terms of bits information gain (x 10^−4^).

## 4. Discussion

Here, we show fMRI evidence that dogs’ brains tended to classify odor mixtures configurally. To test neural mechanisms of dogs’ perception of odors and a mixture, we used fMRI to examine changes in brain activation to previously trained odors associated with reward or no reward, as well as a mixture of the two. Our results suggest that while dogs may have different odor-outcome associations with each individual odor, they perceive the combination of odors as a new odor. In reward processing regions of the brain, we anticipated that if dogs treat mixtures as the sum of their components, then the neural activation to the mixture should be equivalent to the sum of the activation to the reward and no reward components. However, significant differences in activation within the amygdala showed that dogs did not treat these as equivalent conditions.

Further, using machine learning, we identified additional regions of the dog brain, including the peri-amygdalar cortex and the posterior cingulate that significantly predicted the identity of the odor beyond the regions specified in *a priori* hypotheses. Moreover, we found that a multilabel model significantly outperformed a multilabel model, further supporting the conclusion that dogs processed the mixture configurally rather than elementally.

One possible explanation for our results is that the dogs’ perception of odor mixtures may depend on the combined ratio of the odor elements. For example, rabbits trained on a target odor B treated the A+B (ratio 68/32) mixture as elemental but the A+B (ratio 30/70) mixture as configural (Schneider et al., 2016). As our study utilized a 50/50 mixture, we cannot similarly conclude ratio-based differences in elemental or configural processing of odor mixtures in the dog brain. However a second possible explanation is that dogs classify mixtures as themselves as in the 3-way model, but when limited to two classes as in the 2-way model, dogs’ neural biases for novelty influences predictions toward the distractor odor (Prichard, Cook, et al., 2018).

Because the univariate model suggested that dogs treat mixtures as more like the no reward stimulus than the reward stimulus, the mechanism underlying dogs’ discrimination of odor mixtures may have been a learned association between the mixture with absence of reward. The apparent differences in activation in the caudate nucleus and amygdala to odor stimuli associated with reward or no reward suggested that perception changed over time, consistent with a learned discrimination. The significant differential effect for reward versus no-reward across multiple ROIs is therefore consistent with prior research, showing that reward processing regions of the canine brain change in activation relative to the value of conditioned stimuli regardless of modality (Cook et al., 2016; Prichard, Chhibber, et al., 2018). Further, we have previously shown that in an associative reward learning paradigm, changes in the neural activation within the caudate and amygdala within an initial span of 6 minutes, suggesting that a mixture-no reward association could also form quickly (Prichard, Chhibber, et al., 2018). If true, the overall activations in the amygdala and caudate might simply index their relative salience, but not their full identities.

The RFC identified regions important for odor processing similar to those in the human studies, including the amygdala, piriform cortex, and posterior cingulate. In human classical conditioning paradigm using odors, MVPA analyses revealed predictive representations of identity-specific reward in OFC and identity general reward in vmPFC. Reward related functional coupling between OFC and piriform cortex and between vmPFC and amygdala further revealed parallel pathways that support identity-specific and general predictive signaling (Howard et al., 2015; Howard et al., 2016; Zelano, Mohanty, & Gottfried, 2011). Our study also mirrors some of the results examining human’s perception of odor mixtures. In humans, common neural activation patterns in the superior temporal gyrus, caudate nucleus, and insula occur in response to mixtures containing pleasant and unpleasant odors (Bensafi et al., 2012). Given the similar results to human studies, this suggests that shared neural mechanisms may exist across species for odor processing. Further, we show that RFC is a successful classifier for fMRI analyses, with the caution that specific classifiers may be better suited for some studies over others (Misaki, Kim, Bandettini, & Kriegeskorte, 2010).

What does this mean for odor processing in dogs? Understanding how a dog discriminates between odor mixtures can aid in the design of more effective protocols to increase a dog’s performance on odor detection and identification tasks. Protocols designed based on the dogs’ perceptual abilities are less prone to biases inherent to behavioral studies (e.g. the Clever Hans effect) that require human-reported measures. In addition, the dogs’ perception of the mixture stimulus in our study suggests that dogs perceive the mixture as a new odor rather than as its individual elements. Consistent with previous behavioral studies, this may explain why dogs trained on individual target odors have difficulty generalizing to mixtures, but dogs trained on mixtures perform well on detection tasks and detect the target odor when mixed with novel distractors (Hall & Wynne, 2018; Lazarowski & Dorman, 2014; Lazarowski et al., 2015). Further, dogs’ brain activations showed more similarity between the mixture of odors and a no reward odor, suggesting either a learned association or a neural bias toward the no reward odor. Treating a mixture as a novel odor, or having bias toward the no reward component of a mixture, would likely lead to increased false-negatives during a detection task whereas a learned association for mixtures may conflict with detection applications. Knowledge of dogs’ classification of odor mixtures in the dog brain should improve training practices for working dogs and highlight the potential learning aspects inherent in mixture detection tasks. Perceptually driven protocols may therefore enhance a working dog’s detection performance, contributing to the health and safety of humans.

The opportunity to study the neural mechanisms of odor processing in an awake dog also offers two clear advantages over the study of odor processing in humans. First, unlike human studies, dog fMRI offers a unique opportunity to study odor processing in primary sensory areas like the olfactory bulbs given its large size relative to the rest of the dog brain. In humans, the olfactory bulbs is proportionately smaller than in canines, making imaging difficult due to its size and the susceptibility artifact around the sinuses. In our study, the olfactory bulbs were structurally defined in each dog prior to analysis, allowing us to account for the unique aspects of brain morphology across individual canines. In dogs, we found a significant main effect for the differentiation between odor types, similar to human studies of olfactory processing, but over a much larger region of cortex. In other nonhumans, imaging mammalian olfactory cortex may prove difficult due to the resulting signal loss from the air-to-tissue contact in regions near the olfactory bulbs. That said, fMRI of odor processing in canines within this primary sensory region may offer opportunities to understand the mechanisms of odor perception above and beyond what is possible in human fMRI.

Second, while humans use language to describe events and percepts, odors are difficult to describe verbally (Cain, de Wijk, Lulejian, Schiet, & See, 1998; Iatropoulos et al., 2018). When odors are administered during language-dependent tasks, interference occurs when the odor and label are simultaneously processed. This difficulty is thought to be due to limitations in cortical networks, as spatiotemporal patterns produced in neural coding of odors and language are similar. Additionally, humans’ limited language for odors may be a cause for our disregard of this sense compared to our bias for visual stimuli (Lorig, 1999). Odor naming may also account for some of the difficulty reported by participants when attempting to evoke images of the odor objects (Stevenson, Case, & Mahmut, 2007). The inability to name objects based on their olfactory, as opposed to their visual appearance, may be explained by the brain circuitry involved in associating olfactory and visual object features to their lexico-semantic representations (Olofsson & Gottfried, 2015; Olofsson et al., 2014; Olofsson & Wilson, 2018). Dogs prove to be a valuable model for the study of odor processing because they do not have the confound of language and have unique brain morphology for imaging of primary olfactory cortex.

This study also contributes significantly to the existing literature on odor processing in canines. First, this study replicates findings from our previous odor fMRI study using the same dogs and the same odor stimuli (Prichard, Chhibber, et al., 2018). Second, ours is the first study to use data directly from the awake, unanesthetized dog (i.e. brain imaging) as opposed to behavioral outcomes to assess dogs’ perception of odor mixtures. And in contrast, other canine fMRI studies examining the neural correlates of odor processing have used restrained or anesthetized subjects (Jia et al., 2014; Siniscalchi, 2016; Thompkins, Deshpande, Waggoner, & Katz, 2016). Third, we used RFC to perform decoding of the dog brain with awake, unrestrained dogs. In particular, this study supports the differences inherent in univariate fMRI analyses compared to MVPA analyses, as the latter do not classify stimuli based on mean activations (Hebart & Baker, 2018). This allowed us to identify regions supporting classification of stimuli in addition to those specified in univariate analyses. And fourth, ours is the first study to use RFC in nonhuman fMRI and to back map the feature importances into brain space to identify regions that contribute to high classification accuracy. Our novel use of RFC can inform future brain decoding studies as it offers an alternative approach to popular searchlight methods for localizing important regions.

There are several possible limitations to our study. First, the presence of the human owner was constant. Because the human was not blind to the nature of the stimuli, they could have inadvertently influenced the dogs through body language. However, the olfactory stimuli were least likely to be picked up by the humans and were not communicated by human owners, so Clever-Hans effects are unlikely to explain these results. Second, the effects of habituation counteract those of learning. Habituation was perhaps most evident in the amygdala, which displayed a generally declining response with run across trial types. There is ample evidence that the amygdala habituates to repeated presentations of the same stimuli and specifically to odor stimuli (Gottfried, O’Doherty, & Dolan, 2002; Plichta et al., 2014; Poellinger et al., 2001; Wright et al., 2001). It would not be surprising that repeated presentation of the stimuli could lead to decreased physiological response, especially to odors. Third, the pet dogs that participated in the study were not previously trained on odor-detection or discrimination (except for two dogs). Highly trained working dogs may perform differently than pets. However, the results were consistent across dogs that varied in age, breed, and sex, so generalizability to the population is likely. Fourth, the odor training utilized two component odors and one mixture, so the findings may not generalize to all odor mixtures or all mixture concentrations (Schneider et al., 2016). Finally, the stimulus-reward associations were acquired through a passive task in the scanner. No behavioral tests were conducted to test acquisition of the learned associations or to compare to the neural activations. This task design was chosen to minimize any additional training required for the dogs and as a follow-up to our previously published study on odor learning. As in humans, further study may reveal dissociable neural pathways support the associative and perceptual representations of sensory stimuli (Howard et al., 2016).

Our results highlight potential neural mechanisms that underly the perception of odors in dogs. ROI-based analysis highlights the importance of the amygdala for learned associations and that these associations are maintained over time. Machine-learning analysis of dogs’ perception of an odor mixture suggests that dogs perceive odor mixtures as new odors rather than as their individual components. This finding has important implications for the training of odor detection dogs and serves as a potential mechanism underlying dogs’ poor behavioral performance when generalizing from a target odor to mixture. Future decoding studies of the dog brain may allow us to better understand canine perception and highlight potential neural mechanisms for olfactory processing conserved across species.

## 6. Author Contributions

A.P., M.S., and G.B. designed the research; A.P., R.C., K.A. and G.B. collected the data; A.P, M.S. and G.B trained the dogs, A.P., R.C., J.K., and G.B. analyzed data; and A.P., R. C. and G.B. wrote the paper.

## 8. Acknowledgments

Thank you to all of the owners who trained their dogs to participate in fMRI studies: Lorrie Backer, Rebecca Beasley, Emily Chapman, Darlene Coyne, Vicki D’Amico, Diana Delatour, Jessa Fagan, Marianne Ferraro, Anna & Cory Inman, Patricia King, Cecilia Kurland, Claire & Josh Mancebo, Patti Rudi, Cathy Siler, Lisa Tallant, Nicole & Sairina Merino Tsui, Ashwin Sakhardande, & Yusuf Uddin. This manuscript has been released as a preprint at bioRxiv.org (Prichard et al., 2019).

## 9. Data Availability Statement

The raw data supporting the conclusions of this manuscript will be made available by the authors, without undue reservation, to any qualified researcher.

